# Group II metabotropic glutamate receptors modulate sound evoked and spontaneous activity in the mouse inferior colliculus

**DOI:** 10.1101/2020.06.03.130724

**Authors:** Inga Kristaponyte, Nichole L. Beebe, Jesse W. Young, Sharad J. Shanbhag, Brett R. Schofield, Alexander V. Galazyuk

## Abstract

Little is known about the functions of group II metabotropic glutamate receptors (mGluRs2/3) in the inferior colliculus (IC)—a midbrain structure that is a major integration region of the central auditory system. We investigated how these receptors modulate sound-evoked and spontaneous firing in the mouse IC *in vivo*. We first performed immunostaining and tested hearing thresholds to validate VGAT-ChR2 transgenic mice on a mixed CBA/CaJ x C57BL/6J genetic background. Transgenic animals allowed for optogenetic cell type identification. Extracellular single neuron recordings were obtained before and after pharmacological mGluR2/3 activation. We observed increased sound-evoked firing—as assessed by the rate-level functions—in a subset of both GABAergic and non-GABAergic IC neurons following mGluR2/3 pharmacological activation. These neurons also displayed elevated spontaneous excitability and were distributed throughout the IC area tested, suggesting a widespread mGluR2/3 distribution in the mouse IC.

## Introduction

Glutamate is the primary excitatory neurotransmitter in the brain. It binds to both ionotropic and metabotropic glutamate receptors (mGluRs). Ionotropic receptors, which are ligand-gated cation channels, are important for rapid signal transmission, whereas mGluRs act via G-protein coupled second messenger cascades and are relatively slower. In general, mGluRs have been thoroughly investigated due to their potential as therapeutic targets in several psychiatric and neurological disorders (Li et al., 2015; Nicoletti et al., 2015; Chaki, 2017; Yang et al., 2017; Mazzitelli et al., 2018; Crupi et al., 2019). mGluRs are also expressed throughout the auditory system (Lu, 2014; Tang and Lu, 2018) and might be important targets for pharmacological treatment of hearing disorders. However, our knowledge regarding the role of mGluRs in hearing is very limited.

mGluRs are classified into three distinct groups according to sequence homology, second messenger coupling, and agonist selectivity (Nakanishi, 1992; Niswender and Conn, 2010). The objective of this study was to characterize how group II mGluRs, which include subtypes 2 and 3 (mGluRs2/3), modulate spontaneous and sound-evoked firing in the mouse inferior colliculus (IC)—an integrative hub of the central auditory system. Our interest in group II mGluRs arises from recent results showing that behavioral signs of tinnitus in mice were suppressed by intraperitoneal administration of the mGluR2/3 agonist LY354740 (Galazyuk et al., 2019). Furthermore, intravenous LY354740 injection reduced spontaneous activity in the IC. It is possible tinnitus was suppressed because of this reduced spontaneous firing. However, systemic mGluR2/3 activation does not inform us on the origin of the observed effect. Group II mGluRs are expressed in many brain areas (Wright et al., 2001; McOmish et al., 2016), including many auditory structures (Lu, 2014; Tang and Lu, 2018), and the IC receives multiple ascending and descending auditory projections, as well as some non-auditory inputs (Casseday et al., 2002; Gruters and Groh, 2012). Spontaneous activity suppression in the IC could result from mGluR2/3 actions outside the IC itself. Therefore, in the present study, we utilized local mGluR2/3 activation and tested the hypothesis that receptors expressed within the IC exert modulatory actions *in vivo*.

An *in vitro* study by Farazifard and Wu (2010) provides rationale that group II mGluRs are expressed within the IC. Bath application of mGluR2/3 agonist LY379268 reduced the amplitudes of both excitatory and inhibitory postsynaptic currents. Such results suggest that mGluRs2/3 regulate both glutamatergic and GABAergic synaptic transmission in the IC.

Outside the IC, group II mGluR expression has been found in several central and peripheral auditory structures. Below, we summarized the findings from the mammalian animal models. mGluR2 (but not mGluR3) expression was found at the postsynaptic side of the inner hair cell ribbon synapses (Klotz et al., 2019). mGluR2/3 staining was shown throughout the granule cell domain of the cochlear nuclei, as well as in Golgi and unipolar brush cells of the dorsal cochlear nucleus (Petralia et al., 1996; 2000). In the medial nucleus of the trapezoid body, mGluRs2/3 were preferentially localized in astrocytes surrounding the calyces of Held during the early post-natal days, but were found in the presynaptic calyces of Held and postsynaptic principal cells in adults (Elezgarai et al., 2001). In the lateral superior olivary nucleus, mGluR23 labelling was dense in P4 rats but decreased in P18 animals (Nishimaki et al., 2007). Additionally, mGluR2/3 labelling was found in the medial geniculate body of young rats, especially in the marginal zone (Petralia et al., 1996). Finally, concentrated mGluR2 labelling was demonstrated in layer 4 of the primary auditory cortex, with weaker expression evident in layer 5 (Covic and Sherman, 2011; Lee and Sherman, 2012; Venkatadri and Lee, 2014). *In vitro* electrophysiological recordings from various auditory structures shed some light on the physiological mGluR2/3 roles. Group II mGluRs modulate both excitatory (Lee and Sherman, 2012) and inhibitory (Doleviczenyi et al., 2005; Nishimaki et al., 2007; Liu at al., 2014) synaptic transmission. Moreover, these receptors have both pre-synaptic (Doleviczenyi et al., 2005; Nishimaki et al., 2007; Lee and Sherman, 2012; Liu at al., 2014) and post-synaptic (Ene et al., 2003; Irie et al., 2006; Lee and Sherman, 2012) effects.

In the present study, we performed *in vivo* single neuron extracellular recordings before and after pharmacological group II mGluR activation in the mouse IC. We found that mGluRs2/3 modulate sound level processing in a subset of optogenetically identified GABAergic and non-GABAergic cell types. mGluR2/3 activation resulted in steeper rate-level function slopes, lower thresholds, and increased maximum firing rates. Modulatory effects extended to spontaneous activity: post-drug spontaneous firing rates were elevated. These data provide strong evidence that group II mGluRs are expressed in the IC and are involved in sound level processing modulation—an important step towards a better understanding of mGluR roles in the auditory system. Such knowledge may open avenues for possible development of treatment strategies of hearing disorders using mGluR-targeting drugs.

## Materials and methods

### Animals

All procedures were conducted in accordance with the Northeast Ohio Medical University Institutional Animal Care and Use Committee and NIH guidelines. Efforts were made to minimize animal suffering as well as the number of animals used. Animals were housed on a 12-hour day/night cycle with ad libitum access to food and water.

All mice were bred in our animal facility, with breeding pairs purchased from the Jackson Laboratory (Bar Harbor, ME): two males hemizygous for channelrhodopsin-2 in inhibitory neurons (B6.Cg-Tg(Slc32a1-COP4*H134R/EYFP)8Gfng/J, also known as VGAT-ChR2-EYFP; stock # 014548) and six female CBA/CaJ mice (stock # 000656). Neonatal (P0-P3) F1 generation hybrid offspring were phenotyped by briefly placing them under Zeiss AxioImager.M2 microscope (Oberkochen, Germany) and illuminating with a blue *470 nm light. The brains of the offspring carrying the transgene exhibited green fluorescence through the skull. These animals were used in all experiments. Results are described from thirteen transgenic mice of either sex ranging from four to nine months old.

### Surgery

Aseptic techniques were used for all surgical procedures. Surgical anesthesia was induced with isoflurane (3% induction, about 1.5% maintenance in air) and the mouse was placed in a stereotaxic alignment system (David Kopf Instruments, Tujunga, CA; model 1900). The eyes were covered with an ophthalmic lubricating ointment to avoid corneal drying (Puralube Vet Ointment, Dechra, Overland Park, KS). The skull surface was made clear of debris and dry. One scoop of C&B Metabond powder (Parkell, Edgewood, NY; product S399), 6 drops of C&B Metabond Quick Base (product S398) and one drop of C&B Universal catalyst (product S371) were mixed in a ceramic dish with a wooden stick and applied to the skull—with the exception of the area above the right inferior colliculus (IC)—to allow access for all subsequent electrophysiological recordings. A head-post was assembled from a hexagonal stainless-steel standoff (Unicorp, Orange, NJ; product P129M09F16256) and a screw (Bolt Depot, Hingham, MA; product 7662) threaded on one end. The head-post was held vertically and lowered onto the skull (*1.4 mm anterior from bregma) by a custom-made head-post holder, which was attached to the stereotaxic alignment system’s tool holder (David Kopf Instruments, product 1900-54-A). More Metabond mixture was applied around the head-post to secure it. A round craniotomy was made with 1.8 mm diameter trephine drill bit (Fine Science Tools, Foster City, CA; product 18004-18) over the right IC and covered with Kwik-Sil (World Precision Instruments, Sarasota, FL). The surgical site was treated with an antibiotic cream and 0.5% bupivacaine, whereas ketoprofen (5 mg/kg) served as a systemic analgesic. The animal recovered from surgery for at least one day before the start of electrophysiological recordings.

### Experimental design

The specifics of each experimental procedure will be discussed in detail in the following sections. Here, we provide a short overview of the experimental design. After locating a well-isolated single neuron and determining its characteristic frequency (CF; defined as the frequency at which the lowest sound intensity evoked a response), the cell type (GABAergic versus non-GABAergic) was then identified. Then, pre-drug rate-level functions (RLFs) were recorded three or four times. The drug (or the drug vehicle) was then applied topically on the craniotomy. In some experiments iontophoresis was used instead, in order to verify the results obtained using topical application. Post-drug RLFs were then recorded several times. Recording sessions typically continued for 1.5 – 2 h with three sessions conducted on each animal on separate days. At the end of each recording session, the recording site was labelled, then craniotomy rinsed with saline and covered with Kwik-Sil (World Precision Instruments). After the end of the last recording session, mice were sacrificed for immunohistochemistry for the recording site and IC subdivision identification.

### *In vivo* extracellular recordings

Electrophysiological recordings were conducted in a sound-insulated chamber (Industrial Acoustics Company, Bronx, NY). The mouse was briefly anesthetized with 3% isoflurane in air and then administered an intramuscular hind leg injection of a mixture of ketamine (100 mg/kg), xylazine (10 mg/kg), and acepromazine (3 mg/kg). The head-post was secured in a custom-made holder and served as the ground. The eyes were first covered with an ophthalmic lubricating ointment to prevent ocular dryness, and then with small pieces of black light-impermeable material to avoid laser light stimulation. The animal’s temperature was maintained at 37°C with a feedback loop-controlled blanket (Harvard Apparatus, Holliston, MA; model 50-7220F).

Single-unit extracellular spikes were recorded with micropipettes which were pulled from quartz glass tubes (Sutter Instrument, Novato, CA; product QF100-50-10) using a laser electrode puller (Sutter Instrument, model P-2000). Each micropipette tip was broken by a gentle hit against a hanging Kimwipe to create a tip diameter of 1-2 μm. The micropipettes were filled with a dye (see below ‘Recording site labeling’) and 1 M NaCl (electrode resistance was about 15-40 MΩ). Additional control experiments were performed using iontophoresis, which required manufacturing multi-barrel electrodes (see below ‘Manufacturing multi-barrel electrodes’).

The craniotomy was viewed under a stereo microscope (Leica Microsystems, Buffalo Grove, IL; model M80) to remove the dura at the site of the electrode penetration and to position the electrode above the IC using a motorized micromanipulator (Sutter Instrument, model MP-285). The electrode was advanced into the IC in 2 - 4 μm steps by a micropositioner (David Kopf Instruments, model 2660). Extracellular action potentials were amplified (5000X) and bandpass filtered (1000 - 3000 Hz) with an extracellular preamplifier (Dagan Corporation, Minneapolis, MN; model 2400A), then digitized at 40 kHz sampling rate (DataWave Sciworks, Loveland, CO; model DW-USB6432-37-A).

### Acoustic stimulation

Sound intensities were calibrated using custom software written in MATLAB (TytoLogy by S.J. Shanbhag; https://github.com/TytoLogy) and Brüel and Kjær (Duluth, GA) equipment: ¼ inch condenser microphone (model 4939), preamplifier (model 2670), and conditioning amplifier (Nexus model 2690). All sound stimuli were generated at 500 kHz sampling rate and 16-bit depth using BrainWave software (DataWave), attenuated (Tucker-Davis Technologies, Alachua, FL; PA5 programmable attenuator), filtered (Krohn-Hite, Brockton, MA; model 3384), and amplified (Parasound Products, San Francisco, CA; model A23). Sounds were presented free-field, 45° to the left of the mouse mid-sagittal plane and contralateral to the recorded right IC. The distance between the animal and the speaker (LCY-K100 ribbon tweeter; Ying Tai Trading, Hong Kong) was 10 cm.

All tones were 50 ms in duration (5 ms rise/fall times) and were presented at a rate of 4/s, except for the optogenetic cell type identification as described below. Tones used as search stimuli were presented at 70 dB SPL from 6 to 63.3 kHz in 1/4-octave steps. The same tone frequencies were used to determine the CF; sound intensity was reduced in 5 or 10 dB SPL steps. For the RLFs, a tone was presented at the neuron’s CF from 0 to 80 dB SPL, in 5 dB SPL increments. Each intensity was repeated 10 times.

### Pharmacological group II mGluR activation

To pharmacologically activate group II mGluRs, which include subtypes 2 and 3, we used a potent and specific agonist LY354740 (Tocris, Minneapolis, MN; product 3246; Monn et al., 1997; Schoepp et al., 1999). In order to verify the results obtained with LY354740, we utilized another widely used group II mGluR agonist LY379268 (Tocris, product 2453; Monn et al., 1999). Both compounds were prepared in sterile water and pH increased to 8.5 using 1 N NaOH. The drug vehicle was used in control experiments. One μl of drug (or vehicle) solution was applied directly to the craniotomy above the IC using a pipettor. 90 μM LY354740 was used for the initial experiments and later the concentration of the drug was lowered to 30 μM. LY379268 was used at a concentration of 30 μM. These concentrations were selected to avoid possible non-specific drug effects (Schoepp et al., 1999). For iontophoretic application, the concentration of LY354740 was 5 mM as in previous studies (Copeland et al., 2012; 2013; 2017).

### Optogenetic cell type identification

Cell types were determined using a similar method as previously implemented by Ono et al., 2016, 2017. 473 nm laser light (Ready Lasers, Anaheim, CA; MBL-III-473 diode-pumped solid-state laser, fiber collimator, and 400 μm fiber patch cable) was used to illuminate the craniotomy over the IC via a 2 cm long, 400 μm diameter fiber optic cannula (Thorlabs, Newton, NJ; CFMC14L20). The tip of the cannula was positioned about 2 mm above the craniotomy. BrainWave generated a TTL pulse to control the laser output timing. Laser intensity at the tip of the cannula was calibrated with a photodiode power sensor (Thorlabs, S140C) connected to a digital power and energy meter (Thorlabs, PM100D) and then manually adjusted for each experiment depending on the depth of the recording site (intensity varied from about 10 to 45 mW).

When light pulses (30 ms duration, presented every 3 seconds, 15 repetitions) evoked firing, a neuron was classified as GABAergic. If light did not evoke firing, in some neurons further testing was completed to verify that light stimulation indeed reached the area of interest. First, responses to white noise (70 dB SPL, 200 ms duration, presented every 3 seconds, 15 repetitions) were recorded. If a neuron did not respond to white noise, a pure tone at the neuron’s CF was used instead. Then, the same sound protocol was presented together with 30 ms duration light pulse. The light onset was adjusted to occur about 5-10 ms before the neuron’s response onset. The total number of spikes was compared between the sound only and sound with light conditions. Light typically reduced the number of spikes by more than 50%, verifying light stimulation efficiency.

### Manufacturing multi-barrel electrodes and iontophoresis

Single micropipettes were pulled from quartz glass tubes (Sutter Instrument, product QF100-50-10) and then bent (to an angle of about 40°, at approximately 5 mm away from the tip) using a laser electrode puller (Sutter Instrument, model P-2000; part FPS is a special P-2000 accessory for bending the tips). The micropipette tips were gently struck against a hanging Kimwipe, breaking-off to create a tip diameter of about 1-2 μm.

3-barrel micropipettes were prepared by placing about 1 cm length, 4 mm diameter polyolefin heat shrink tubing (Innhome, Ontario, CA; product HS-532B) about 1 cm away from the ends of 3-barrel borosilicate glass capillary tubes (Sutter Instrument, product FG-G3BF100-75-10) and shrinking it with an infrared lamp. The 3-barrel micropipettes were then pulled on a Gravipull-3 micropipette puller (Kation Scientific, Minneapolis, MN) and broken to an overall tip diameter of about 5 μm using a previously described method (Dondzillo et al., 2013).

Single and 3-barrel micropipettes were aligned and then glued together in a piggyback configuration (Havey and Caspary, 1980) by first applying black super glue (StewMac, Athens, OH) and then 5 Minute Epoxy (Devcon, Hartford, CT) using a similar method as described by Dondzillo et al. (2013).

Iontophoretic micropipettes were connected to Dagan 6400 advanced micro-iontophoresis current generator which provided independent control of two iontophoresis channels (drug/vehicle) and a balancing channel. Current parameters were selected based on previous studies which utilized LY354740 iontophoresis (Copeland et al., 2012; 2013; 2017): positive 15 nA retaining current prevented spontaneous drug diffusion, whereas ejection current was negative 25 nA.

### Recording site labeling

After neuronal activity was assessed, recording sites were labeled with either 1% Neurobiotin 350 in 1 M NaCl (Vector laboratories, Burlingame, CA; product SP-1155), 1% Neurobiotin 488 in 1 M NaCl (Vector laboratories, product SP-1125-2), or 0.1 % Fluoro-Gold in saline (Fluorochrome, Denver, CO). Different dye colors distinguished the recording sites labelled on different recording days in the same animal. When a single micropipette was used, a drop of dye solution was placed on its back end and allowed to move to the tip; the micropipette was then filled with 1 M NaCl solution. All dyes were ejected using Dagan 6400 advanced micro-iontophoresis current generator (the wire which connected the micropipette to the extracellular preamplifier during recordings was switched to the wire connected to Dagan 6400). Positive 1000 nA current was used to eject Neurobiotin 350 and Neurobiotin 488 (2 and 4 min, respectively) and negative 1000 nA current for Fluoro-Gold (4 min). The magnitude of the current is at least 10 times smaller than those used to cause electrolytic lesions (10 - 50 μA; Townsend et al., 2002; Ayala and Malmierca, 2015; Harris et al., 2017; Yang et al., 2020). Indeed, our histological examination did not reveal any lesions at the recording sites.

### Recording site identification

After the end of the last recording session, mice were injected with an overdose of Fatal-Plus (>100 mg/kg, i.p., Vortech, Dearborn, MI). Following loss of corneal and withdrawal reflexes, the animal was decapitated, the skull partially removed, and the head placed in 4% paraformaldehyde (PFA) at room temperature for 30-60 min. Then, the brain was removed and stored at 4°C in 4% PFA containing 25% sucrose overnight. The midbrain was trimmed out, frozen, and coronal sections were collected from the IC on a sliding microtome at a thickness of 40 or 50 μm. IC sections were mounted serially from a 0.2% gelatin solution onto gelatin-coated slides.

Slides were air-dried overnight, then sections were stained on-slide for GAD67 and GlyT2 to determine IC subdivisions (Buentello et al., 2015). A hydrophobic barrier pen was used to outline all sections on the slide. Once the hydrophobic barrier was dry, slides were rinsed in phosphate-buffered saline solution (PBS, 0.9% NaCl in 0.01 M phosphate buffer (PB), pH 7.4), then permeabilized in 0.3% Triton X-100 in PBS for 30 minutes at room temperature. A blocking solution made up of 0.1% Triton X-100 and 10% normal goat serum (NGS) in PBS was applied for one hour at room temperature. The primary antibody solution containing 1% NGS, 0.1% Triton X-100, mouse anti-GAD67 (Millipore Sigma, St. Louis, MO; MAB5406; 1:250) and guinea pig anti-GlyT2 (Synaptic Systems, Goettingen, Germany; 272-004; 1:2500) was applied to the slides, and slides were placed in a sealed container overnight at 4°C. The next day, slides were rinsed in PBS, then a secondary solution containing Alexa Fluor 750-labeled goat anti-mouse (Molecular Probes, Eugene, OR; A21037; 1:100) and Alexa Fluor 647-labeled goat anti-guinea pig (Molecular Probes, A21450; 1:100) in PBS was applied for one hour at room temperature. Slides were rinsed in PBS, air dried, and coverslipped with DPX mounting medium (Sigma-Aldrich, St. Louis, MO).

Slides were examined on a Zeiss AxioImager.Z2 microscope attached to a Neurolucida system (MBF Bioscience, Williston, VT). Images were taken with a 5X objective. In sections where a labeled recording site could be identified, the section was outlined using the Neurolucida system, subdivision outlines were added based on the GAD67 and GlyT2 staining (Buentello et al., 2015), and the recording site was marked. The outline was exported, then outlines from all cases with identifiable recording sites were overlaid onto a representative series in Adobe Illustrator.

### Transgenic mouse validation

To test whether F1 hybrid offspring between VGAT-ChR2-EYFP and CBA/CaJ mice correctly expressed channelrhodopsin-2 in inhibitory neurons, two four-month-old mice (one male and one female) from different breeder pairs were sacrificed for immunostaining. Each animal was deeply anesthetized with an overdose of Fatal-Plus (>100 mg/kg, i.p., Vortech) and perfused transcardially with 0.1 M PB to clear the blood, then with 50 ml of 4% paraformaldehyde in 0.1 M PB, followed by 50 ml of the same fixative containing 10% sucrose. Brains were removed and post-fixed in fixative containing 25% sucrose overnight. The following day, the midbrain was trimmed out, frozen, and coronal sections were collected from the IC on a sliding microtome at a thickness of 40 μm. Sections were collected in three series. For one series, free-floating sections were stained for GAD67 to confirm that EYFP-expressing cells in the IC were indeed GABAergic. Sections were rinsed in PBS, then permeabilized with 0.2% Triton X-100 in PBS for 30 minutes at room temperature. Nonspecific staining was blocked by incubating sections in a solution containing 10% normal goat serum (NGS) and 0.1% Triton X-100 in PBS for one hour at room temperature. A primary antibody solution containing 1% NGS and 0.2% Triton X-100 in PBS was applied overnight at 4°C. Primary antibodies used were anti-GAD67 (Millipore Sigma, MAB5406; 1:250; to label GABAergic cells) and anti-GFP (Thermo Fisher Scientific, Waltham, MA; A10262; 1:400; to enhance EYFP fluorescence-the structure of GFP and EYFP are similar enough that anti-GFP antibodies can recognize EYFP). The next day, slides were rinsed in PBS, then a secondary solution containing Alexa Fluor 488-labeled goat anti-chicken (Molecular Probes, A11039; 1:100) and Alexa Fluor 546-labeled donkey anti-mouse (Molecular Probes, A10036; 1:100) in PBS was applied for one hour at room temperature. Sections were rinsed in PBS, then mounted from a 0.2% gelatin solution onto gelatin-coated slides. Slides were air dried and then coverslipped with DPX mounting medium (Sigma-Aldrich).

Photomicrographs were taken using a Zeiss AxioImager.Z2 microscope with a 63X oil-immersion objective (NA = 1.4) and an Apotome 2 to provide optical sectioning at 0.5 μm depth intervals. The high magnification images shown below are maximum intensity projections of collected stacks. Adobe Photoshop was used to crop and colorize images, and globally adjust levels when necessary.

### Auditory brainstem response (ABR) testing

ABR recordings were obtained under ketamine/xylazine anesthesia (100 mg/kg and 10 mg/kg, respectively). Stainless-steel electrodes (LifeSync Neuro, Coral Springs, FL; product S02918-B) were placed subdermally, one at the ventral edge of each pinna, one along the vertex and the fourth one at the tail. The speaker (LCY K-100 Ribbon Tweeter) was placed 10 cm from the animal’s head. Custom OpenEx software controlling the TDT processor (Alachua, FL; model RZ6) generated all stimuli and recorded responses. Tone pips were 5 ms duration (with 0.5 ms rise/fall time), at 4, 8, 12.5, 16, 20, 25, and 31.5 kHz, from 75 to 5 dB SPL (5 dB SPL steps, 300 repetitions at each sound level), and presented at 50 Hz. ABR threshold was considered the lowest sound level at which a waveform could be observed.

### Electrophysiological data analysis and statistics

Electrophysiological data was first processed using BrainWave software (DataWave). For all recordings obtained from an individual neuron, spikes were extracted using a single visually determined threshold. To confirm a well-isolated single neuron recording, all spike waveforms were superimposed for visual examination. Spike data were binned (1 ms bins) and exported for all further analyses using R (R Core Team, 2020).

Rate-level functions were fit using a non-linear five parameter logistic model (Watkins and Barbour, 2011a; Watkins and Barbour, 2011b; R script provided in the Appendix). Three values were extracted from each fitted curve: maximum firing rate, threshold, and saturation. Threshold and saturation values corresponded to sound level at 20% and 80% of the maximum firing rate, respectively. The slope was calculated as a ratio between the firing rate’s dynamic range and the range of sound levels between the threshold and saturation points.

Previous electrophysiological studies typically observed that mGluR2/3 pharmacological targeting had effect in only a subset of neurons tested (Sanes et al., 1998; Voytenko and Galazyuk, 2011; Galazyuk et al., 2019). This could be due to several reasons, such as lack of uniform receptor expression on all neurons or incomplete receptor activation because of limited drug concentration at the recording site. Thus, we used k-means cluster analysis to assign neurons to either drug effect or no drug effect groups.

When raw data was not normally distributed, we used the Box-Cox procedure, which provides an optimal transformation to normality for non-normally distributed samples (Box and Cox, 1964). In a case where transformed data were still not normally distributed (as evaluated semi-qualitatively using Q-Q plots; Sokal and Rohlf, 2011), statistical analyses were performed using rank-transformed data (Conover and Iman, 1981). The mixed-effects statistical approach (Pinheiro and Bates, 2000) was utilized to account for random variation among individual neurons for population analyses (neuron was treated as a random factor). Non-parametric two-sample Kolmogorov-Smirnov tests were used for comparing monotonicity index distributions before and after drug application. Fisher’s Exact Test was used to assess the difference in proportions between nominal variables (e.g. cell type, sex). Paired t-tests were used to assess within-neuron differences pre- and post-drug conditions. When multiple comparisons were performed, p-values were adjusted using the Benjamini and Hochberg’s false discovery rate procedure (Benjamini and Hochberg, 1995); to indicate those cases, p-values are noted as adj. p in the Results.

To accomplish all analyses, several add-on R packages were required: car (Fox et al., 2011), lme4 (Bates et al., 2015), lmerTest (Kuznetsova et al., 2017), emmeans (Lenth, 2017), plyr (Wickham, 2016), nplr (Commo and Bot, 2016), minpack.lm (Elzhov et al., 2016), ggplot2 (Wickham, 2020), cluster (Maechler et al., 2019), factoextra (Kassambara and Mundt, 2019), knitr (xie, 2020).

## Results

### Channelrhodopsin-2 is correctly expressed in inhibitory neurons of mixed genetic background mice

Zhao et al. (2011) established the VGAT-ChR2-EYFP transgenic mouse line, which expresses channelrhodopsin-2 (ChR2) fused with an enhanced yellow fluorescent protein (EYFP) under the control of the vesicular γ-aminobutyric acid (GABA) transporter (VGAT) promoter. VGAT is expressed in GABAergic and glycinergic neurons and is responsible for loading GABA and glycine from neuronal cytoplasm into synaptic vesicles (Sagné et al., 1997; Gasnier, 2000). We wanted to take advantage of this cell-type specific transgenic mouse line to optogenetically determine whether the recorded neurons are GABAergic or not.

However, this mouse line was developed on the commonly used C57BL/6J strain background, which is not ideal for hearing research aimed at understanding normal auditory function. These animals have early onset hearing deficits due to homozygosity for the *Cdh23^753A^* point mutation, also known as *ahl* mutation (Zheng et al., 1999; Noben-Trauth et al., 2003; Ohlemiller et al., 2016). To avoid hearing abnormalities found in C57BL/6J background, we crossed VGAT-ChR2-EYFP animals with CBA/CaJ mice and used F1 generation hybrid offspring for all experiments. CBA/CaJ mice are homozygous for the wild-type *Cdh23* gene and maintain normal hearing through their lifespan. A single copy of the wild type *Cdh23* allele is sufficient for normal auditory processing until at least one year of age (Frisina et al., 2011; Burghard et al., 2019; Lyngholm and Sakata, 2019).

Because our study is the first to cross VGAT-ChR2-EYFP mice with CBA/CaJ mice, we performed immunostaining to verify that ChR2-EYFP transgene is correctly expressed in inhibitory neurons of mixed genetic background mice. ***Figure 1*** shows representative images of inferior colliculus (IC) central nucleus cells double-labelled with anti-GAD67 and anti-GFP antibodies, which are markers for GABAergic neurons and ChR2, respectively. Consistent with previous studies, we found that neurons that expressed EYFP routinely co-expressed GAD67 in cell bodies (Zhao et al., 2011; Ono et al., 2016; Naumov et al., 2019). We also observed punctate label, probably representative of EYFP-expressing axon terminals, which were also stained for GAD67. Our results confirmed that the ChR2-EYFP transgene is correctly expressed in mixed CBA/CaJ x C57BL/6J genetic background mice, which lack the hearing abnormalities found in the C57BL/6J background.

**Figure 1.**
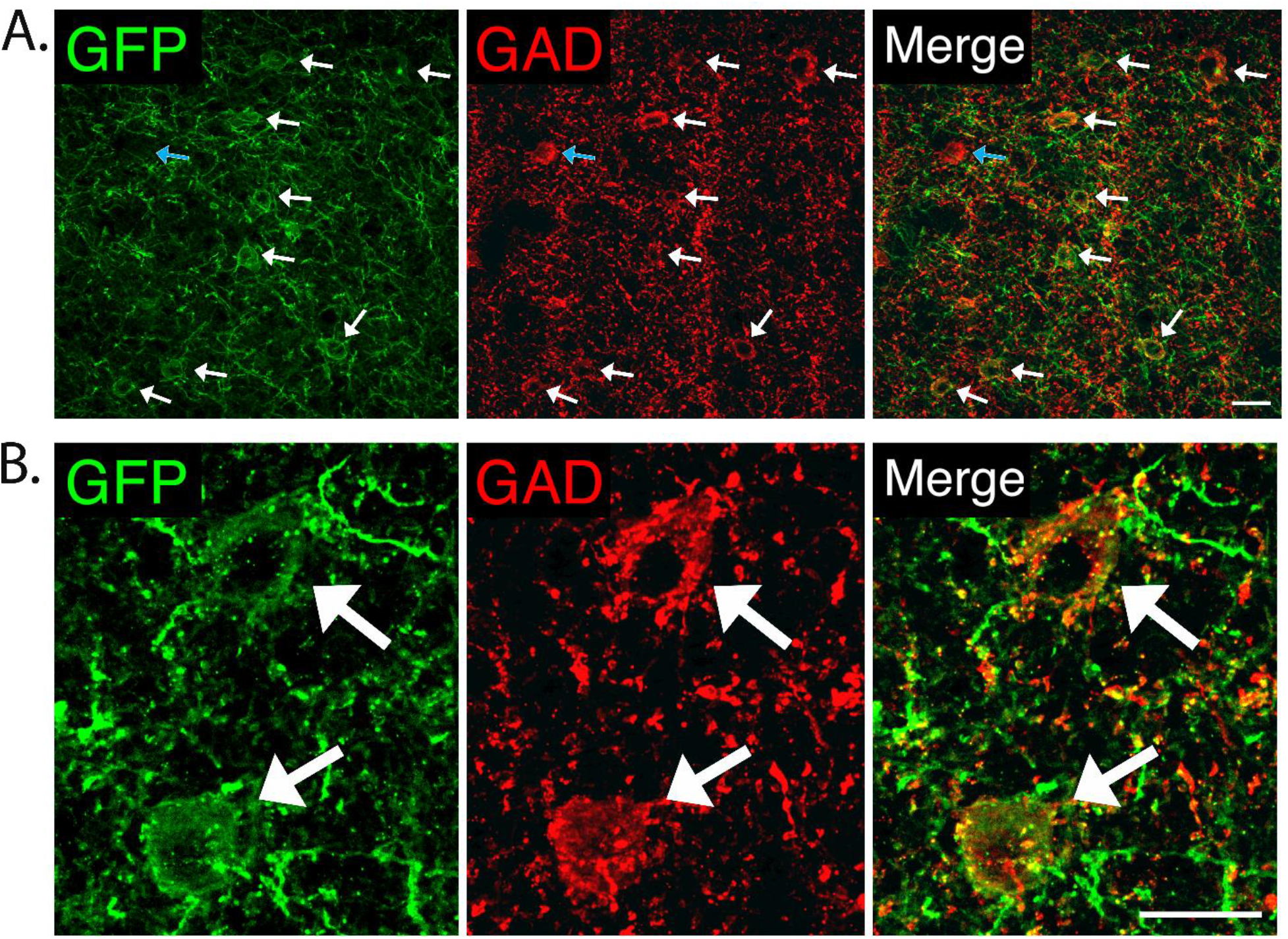
Channelrhodopsin-2 expression in inhibitory inferior colliculus central nucleus neurons of mixed CBA/CaJ x C57BL/6J genetic background mice. (**A**) A low magnification image shows that the majority of inhibitory cells labeled with anti-GAD67 antibody (red) also express ChR2-EYFP labelled with anti-GFP antibody (green). White arrows show examples of double-labelled cells. The blue arrow shows a rare example of a cell that labeled with the anti-GAD antibody but did not express EYFP. Scale = 20 μm. (**B**) High magnification images of double-labelled cells. Scale = 20 μm.

### Mixed genetic background VGAT-ChR2-EYFP mice have normal hearing

To confirm that ChR2-EYFP transgene does not interfere with normal auditory processing, we compared auditory brainstem response (ABR) thresholds between three ChR2-EYFP-positive and three ChR2-EYFP-negative littermates from two different breeder pairs (***Figure 2***). At the time of ABR testing, the animals were about 7.5 months old. For statistical analysis, hearing thresholds were transformed using the Box-Cox procedure to improve normality (Box and Cox, 1964). A mixed-effects ANOVA indicated a significant independent main effect for the stimulus frequency (F_6,66_ = 115.6, p < .0001), whereas the main effect of the genotype, as well as the interaction between the frequency and the genotype, were not significant (F_1, 4_ = .002, p = 0.97, F_6, 66_ = .58, p = 0.75, respectively). Post-hoc pairwise comparisons of estimated marginal means (emmeans), also known as least squares means, showed no significant differences in hearing thresholds between ChR2-EYFP-positive and ChR2-EYFP-negative mice at all frequencies tested (all adj .p > .22). These data demonstrate that at least until 7.5 months of age, F1 generation hybrid offspring from VGAT-ChR2-EYFP and CBA/CaJ mice have normal hearing thresholds.

**Figure 2.**
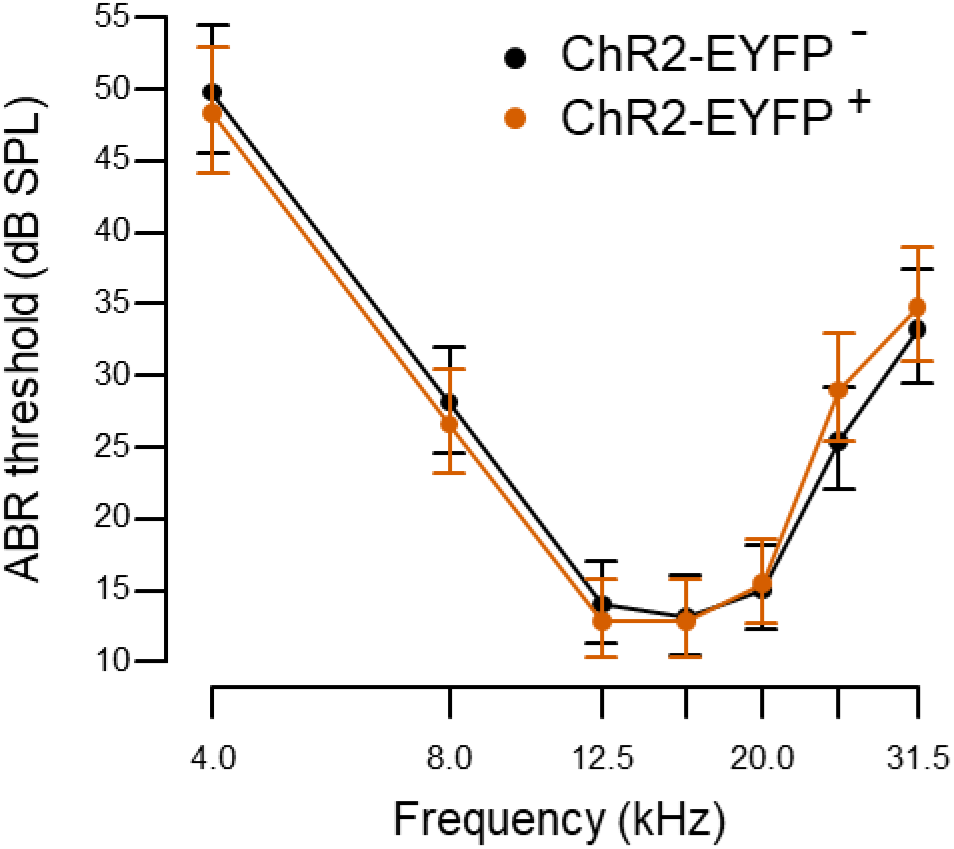
Auditory brainstem response thresholds (expressed as back-transformed least squares means) were not significantly different between ChR2-EYFP-positive (ChR2-EYFP^+^) and ChR2-EYFP-negative (ChR2-EYFP^-^) littermates. Error bars represent 95% confidence intervals of means.

### MGluR2/3 activation enhances sound level processing in a subset of IC neurons

The effect of group II mGluR activation on sound level processing was tested in 31 IC neurons from thirteen VGAT-ChR2-EYFP mixed genetic background mice. Extracellular single-cell recordings were obtained in response to 50 ms duration pure tones presented at the neuron’s characteristic frequency from 0 to 80 dB SPL. Each neuron was tested with the same sound stimulation protocol several times before and after topical mGluR2/3 agonist LY354740 application, resulting in multiple rate-level functions (RLFs). The same analysis window was used to assess all pre- and post-drug data from an individual neuron. The window was adjusted for each neuron to capture sound-evoked spikes based on visual inspection of peri-stimulus time histograms plotted for each RLF.

Each RLF was fit with a non-linear five parameter logistic model and the peak firing rate, slope, as well as threshold were extracted from the fitted curve (***Figure 3***). This RLF quantification approach has been previously used by several research groups (Watkins and Barbour, 2011a; Watkins and Barbour, 2011b; Moore and Wehr, 2013; Stefanescu et al., 2015; Chambers et al., 2016; Ono et al., 2017). To assess how well each curve fit the RLF data, all fits were visually examined. Additionally, we obtained the average error between the fitted curve and the data points, i.e. the residual standard error (RSE). In the final data set, the median RSE was 1.70 and the mode was 0.11. Several RLFs with RSE values larger than ten were excluded from further analyses; this cutoff point was determined by visual inspection of the fits.

**Figure 3.**
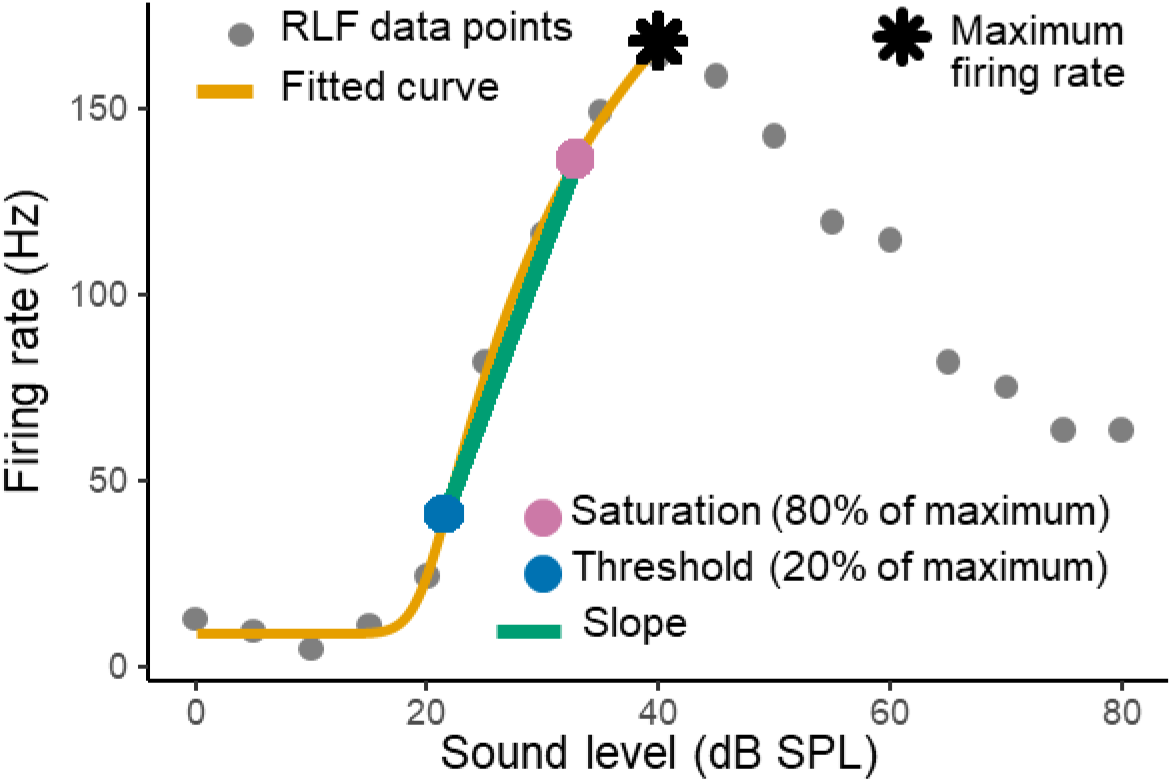
Example rate-level function fitted with a curve. The maximum firing rates, slopes, and thresholds were extracted from the fitted curves for subsequent data analyses.

Each dependent variable (i.e., the peak firing rate, slope, and threshold) was first normalized using the Box-Cox procedure (Box and Cox, 1964). Then, a separate mixed-effects ANOVA test was used for each dependent variable to compare pre-drug and post-drug RLF data obtained from all 31 neurons. After mGluR2/3 agonist application the peak firing rates were significantly higher (F_1,255.1_ = 47.9, adj. p < .0001), the slopes steeper (F_1,255.4_ = 13.8, adj. p < .001), and thresholds became lower (F_1,255.2_ = 38.0, adj. p < .0001). In ***Figure 4*** we plotted mean pre-drug and mean post-drug peak firing rate, slope, and threshold for each neuron, as well as population data visualized with boxplots (the plot presents Box-Cox transformed data).

**Figure 4.**
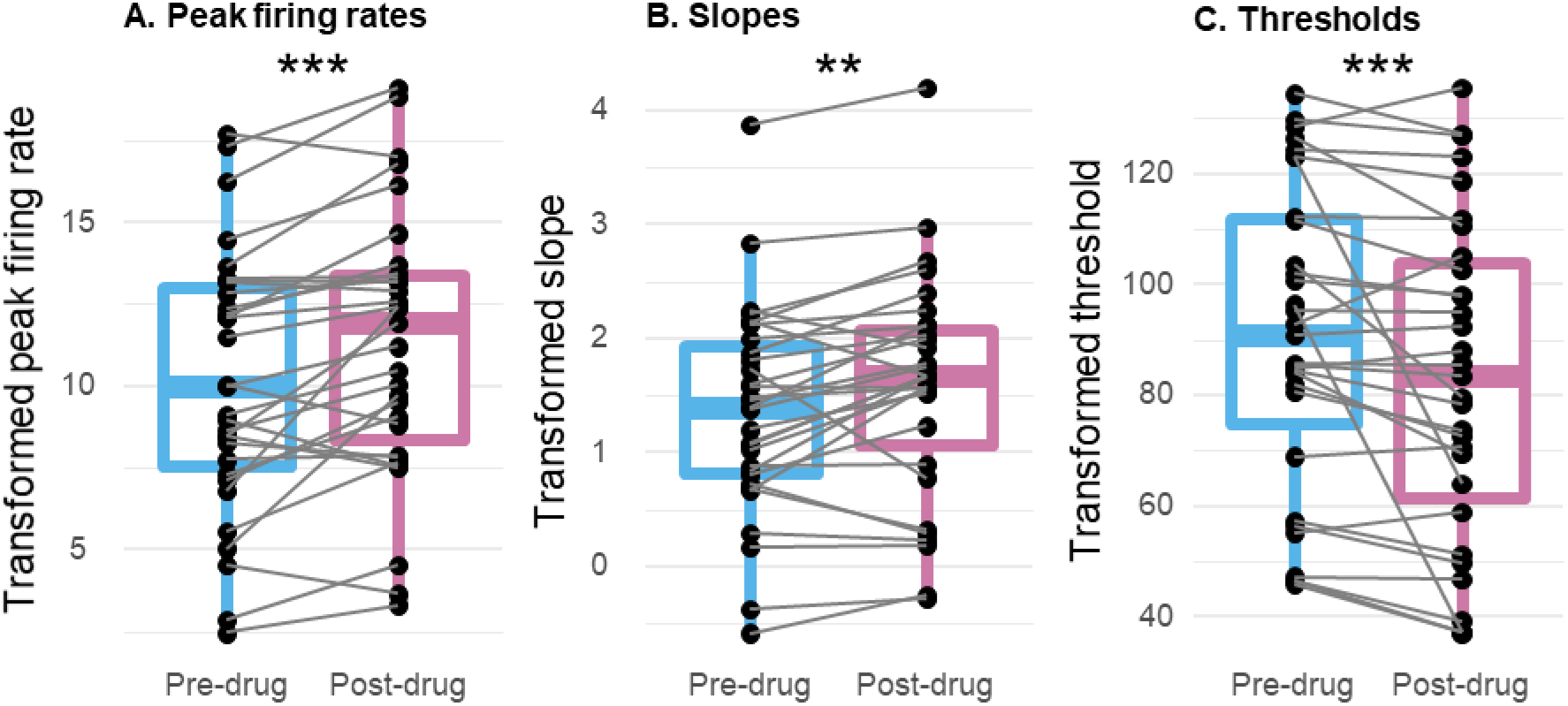
The effect of topical mGluR2/3 agonist LY354740 application on rate-level functions’ peak firing rates (**A**), slopes (**B**), and thresholds (**C**). MGluR2/3 activation significantly increased the peak firing rates, steepened the slopes, and lowered the thresholds. Each neuron’s (n = 31) mean pre-drug and mean post-drug values are connected by grey lines; population data is summarized with boxplots (the plot presents Box-Cox transformed data). **p<0.001, ***p<0.0001.

Even though population-level analysis demonstrated significant RLF changes after mGluR2/3 activation, visual examination of RLFs indicated that the magnitude of drug effect slightly varied between neurons and that the effect was absent in some neurons. Previous studies also described that only a subset of neurons was affected by mGluR2/3 pharmacological manipulation (Sanes et al., 1998; Voytenko and Galazyuk, 2011; Galazyuk et al., 2019). Thus, we utilized cluster analysis to empirically sort the neurons into groups based on the overall similarity of LY354740 effect on their RLFs. Using the differences between pre-drug and post-drug mean peak firing rate and firing rate’s dynamic range as variables for k-means clustering, we found three distinct clusters with eight, eleven, and twelve neurons in each. Empirically derived cluster group was considered a fixed factor in subsequent mixed-effects ANOVA analyses, which were conducted separately on all three dependent variables described in ***Figure 4***, i.e. the peak firing rate, slope, and threshold. Detailed results are provided in ***Table 1***. Post-hoc pairwise comparisons of estimated marginal means showed that each dependent variable was significantly affected by drug application in two clusters, but not in a third cluster. Therefore, for all subsequent analyses we collapsed the first two cluster groups into a single group. ***Figure 5*** shows drug effect on the resulting two groups of neurons. In the first group (***Figure 5A***), mGluR2/3 agonist significantly increased the peak firing rates (F1,169 = 99.0, adj. p < .0001), steepened the slopes (F1,169 = 25.6, adj. p < .0001), and reduced the thresholds (F1,169 = 41.1, adj. p < .0001). No such significant changes were found in the second group [***Figure 5B***; peak firing rates (F_1,85_ = 5.2, adj. p = .08), slopes (F_1,85_ = 0.1, adj. p = .71), and thresholds (F_1,85_ = 1.0, adj. p = .50)]. Thus, we refer to these groups as *drug effect* versus *no drug effect* in subsequent descriptions.

**Table 1.**
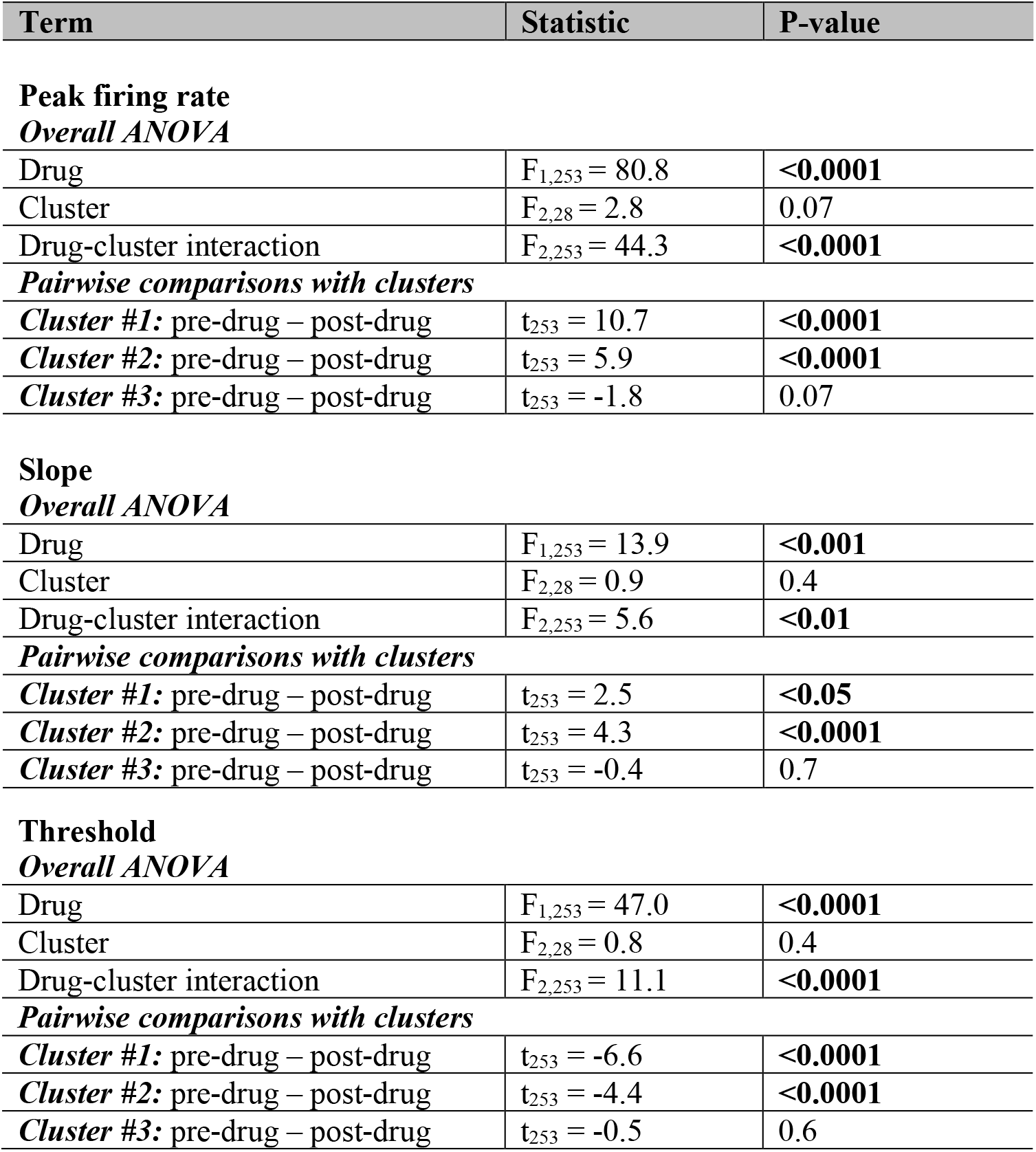
The effect of topical LY354740 application on RLFs’ peak firing rates, slopes, and thresholds. The drug had a significant effect on all three dependent variables in cluster #1 and #2, but not in cluster #3 neurons.

**Figure 5.**
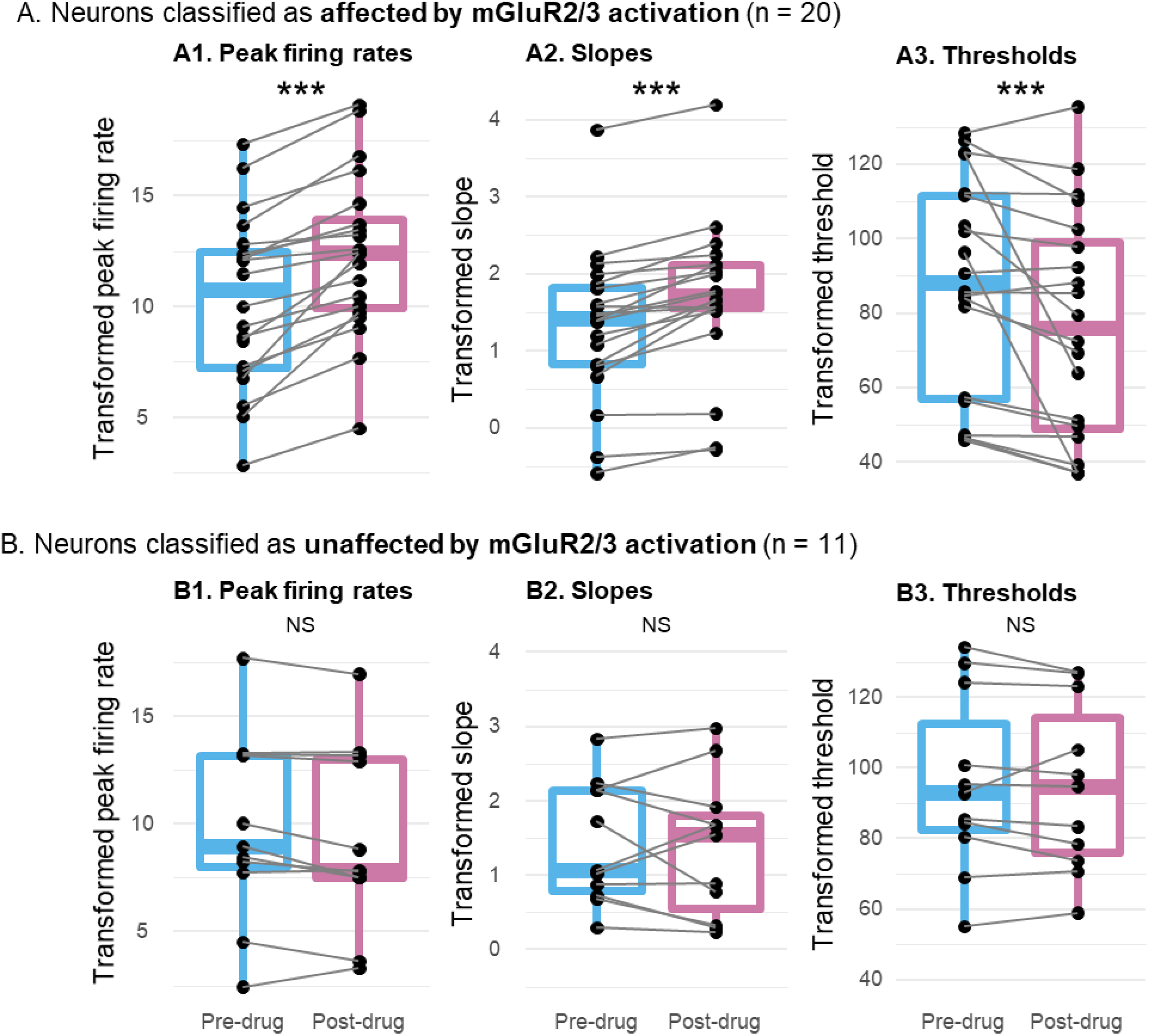
Neurons were assigned to two groups: those which had rate-level functions affected by topical mGluR2/3 agonist LY354740 (**A**; n = 20), and those in which mGluR2/3 activation did not have a clear effect (**B**; n = 11). Box-Cox transformed rate-level functions’ peak firing rates (**A1, B1**), slopes (**A2, B2**), and thresholds (**A3, B3**). Each neuron’s (n = 31) pre-drug and post-drug means are connected by grey lines; population data is summarized with boxplots. ***p<0.0001, NS p>0.05.

We observed that RLFs became slightly more monotonic after drug application. To quantify that, for each RLF we calculated the monotonicity index (MI; Ono et al., 2017): firing rate at the maximum sound intensity level (80 dB SPL) was divided by the maximum firing rate. Neurons with more monotonic RLFs had MIs close to one, as their maximum firing rates occurred at/near the maximum sound intensity level. In ***Figure 6*** we plotted MIs separately for neurons assigned to the drug effect and no drug effect groups. Two-sample Kolmogorov-Smirnov test indicated statistically significant difference between predrug versus post-drug MI distributions in the drug effect group (D = 0.25, p < 0.05), whereas no such difference was found in the no drug effect group (D = 0.16, p = 0.62).

**Figure 6.**
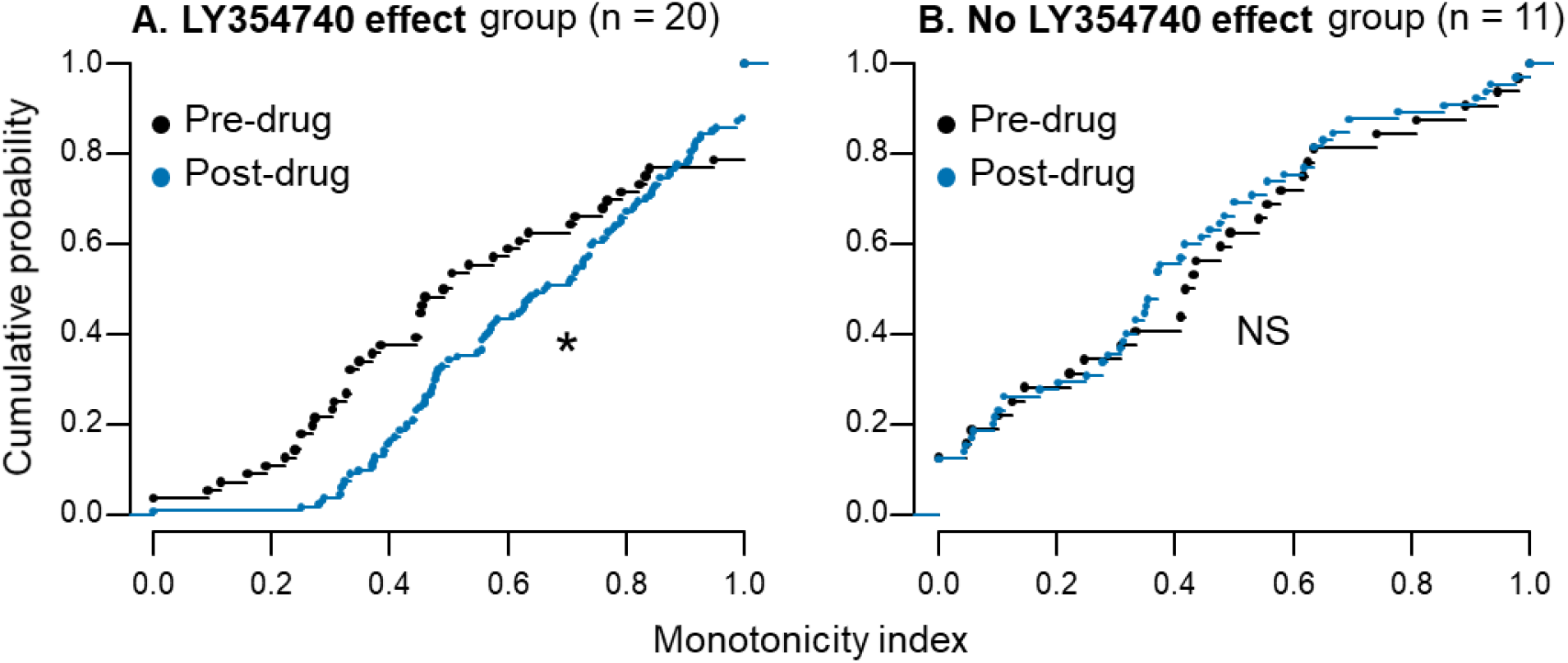
The effect of topical mGluR2/3 agonist LY354740 application on neurons’ monotonicity. All pre-drug and post-drug monotonicity indexes are plotted as cumulative distributions. (**A**) Neurons from the drug effect group (n = 20) became significantly more monotonic after drug application. (**B**) Monotonicity did not significantly change in neurons assigned to the no drug effect group (n = 11). *p<0.05, NS p>0.05.

### MGluR2/3 activation has similar effect on GABAergic and non-GABAergic IC neurons

As experiments were conducted on VGAT-ChR2-EYFP transgenic mice in which inhibitory neurons express channelrhodopsin-2, laser light pulses evoked firing only in inhibitory cells, allowing us to distinguish GABAergic from non-GABAergic cell types optogenetically. From a total of 31 neurons tested with topical LY354740 application, six were inhibitory. Five of them were from the group where mGluR2/3 activation resulted in significant RLF changes (***Figure 5A***). In ***Figure 7*** we plotted example RLFs from both cell types before and after topical LY354740 application. Both monotonic and non-monotonic neurons were affected by the drug. The effect typically occurred at 1 to 5 minutes after drug application and lasted throughout the recording, about 12 to 15 minutes after drug application.

**Figure 7.**
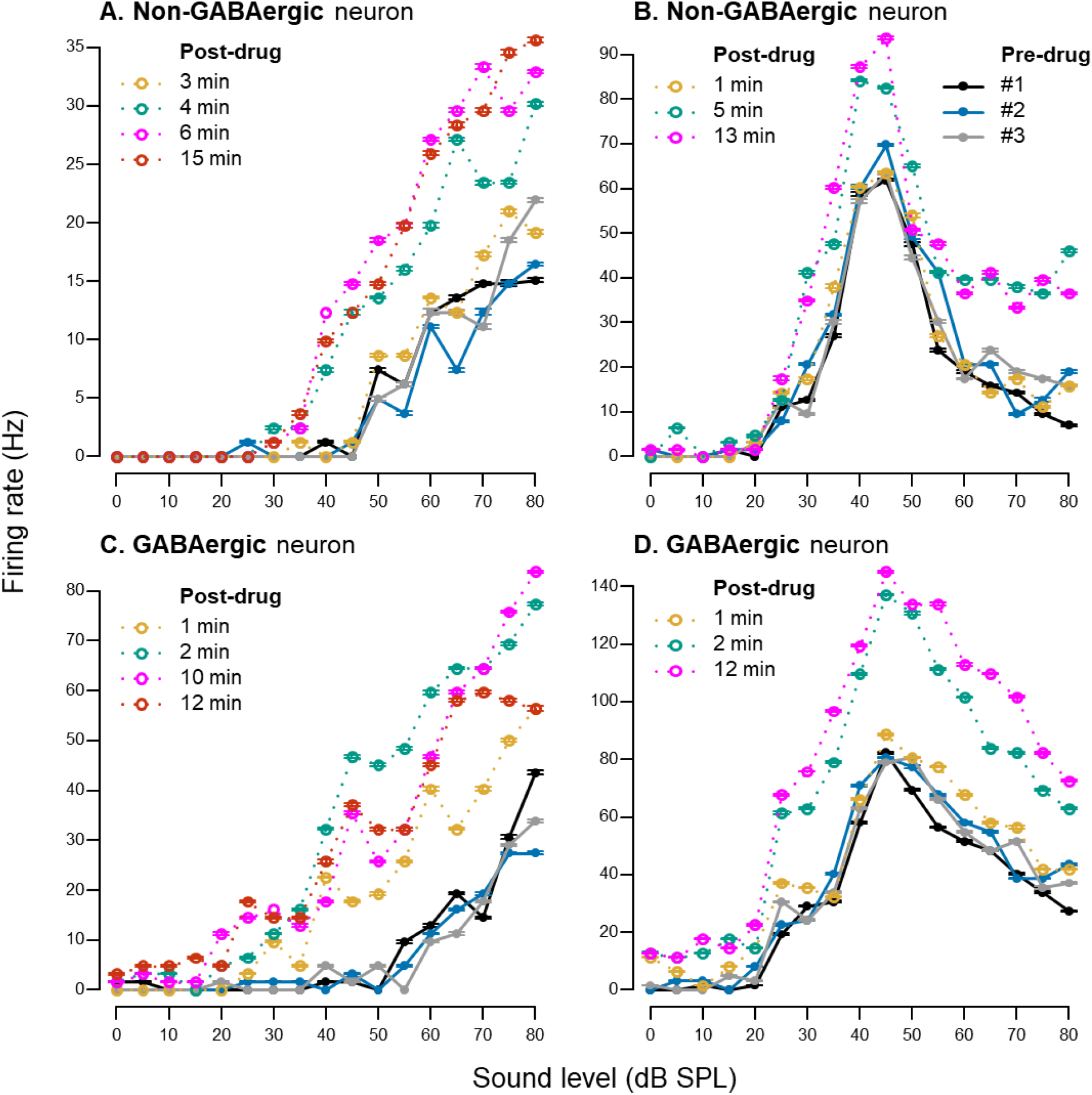
Examples of rate-level functions before and after topical LY354740 application. (**A, B**) Non-GABAergic neurons. (**C, D**) GABAergic neurons. Pre-drug rate-level functions are plotted in solid black, blue, and grey lines in all panels (legend only shown in panel **B**). Post-drug rate-level functions are color coded yellow, green, pink and orange to demonstrate drug effect over time; time after drug application is indicated for each individual rate-level function. Error bars represent ± SEM of ten sound stimulus presentations at each sound level.

### Relationship between mGluR2/3 activation and neurons’ temporal response type

To gain some insight into which IC cell types might be modulated by group II mGluRs, we examined whether mGluR2/3 activation induced RLF changes more often in neurons with certain temporal response type. Also, we asked whether mGluR2/3 agonist changed those temporal response patterns. To answer these questions, for each neuron two peri-stimulus time histograms (PSTHs) were created: one from all pre-drug data, the other from post-drug data. All PSTHs were visually examined and assigned to either phasic/onset or tonic/sustained temporal response pattern. Whereas all neurons maintained their characteristic response pattern after drug application, tonic/sustained type was more common amongst the neurons assigned to the drug effect group. Fourteen of twenty (70%) neurons from this group had tonic/sustained firing. Neurons from no drug effect group displayed almost equal numbers of temporal response patters: 45.5% had phasic/onset firing and 54.5% had tonic/sustained pattern. However, Fisher’s Exact Test did not result in statistically significant association between the temporal response type and drug effect group (p = .45).

### Age, sex and drug dose do not mediate RLF changes in response to mGluR2/3 activation

It has been shown that mGluR expression is developmentally regulated in several brain regions (Catania et al., 1994; Defagot et al., 2002; Doherty et al., 2004; Frank et al., 2011; McOmish et al., 2016; Hernandez et al., 2018). Thus, we examined whether age differences between the animals could explain the variability in LY354740 effect (***Figure 8A***). Two sample t-test demonstrated that there was no significant age difference between the neuron group in which the drug significantly changed the RLFs (mean = 194 days, SD = 47.3 days) versus the group without such changes (mean = 197 days, SD = 43.0 days; t22 = 0.2, p = .86). Furthermore, the animals’ sex also did not explain drug effect variability (***Figure 8B***). Sixteen out of 31 neurons were recorded from female mice and 15 neurons from male mice. According to the Fisher’s Exact Test result, the proportion of male and female mice did not differ by drug effect group (p = .14). Similarly, Fisher’s Exact Test showed that proportion of neurons tested with lower (30 μM) and higher (90 μM) LY354740 concentration was not significantly different between drug effect groups (p = .72; ***Figure 8C***). These data suggest that factors other than sex, age and drug dose determine whether RLF is affected by group II mGluR activation.

**Figure 8.**
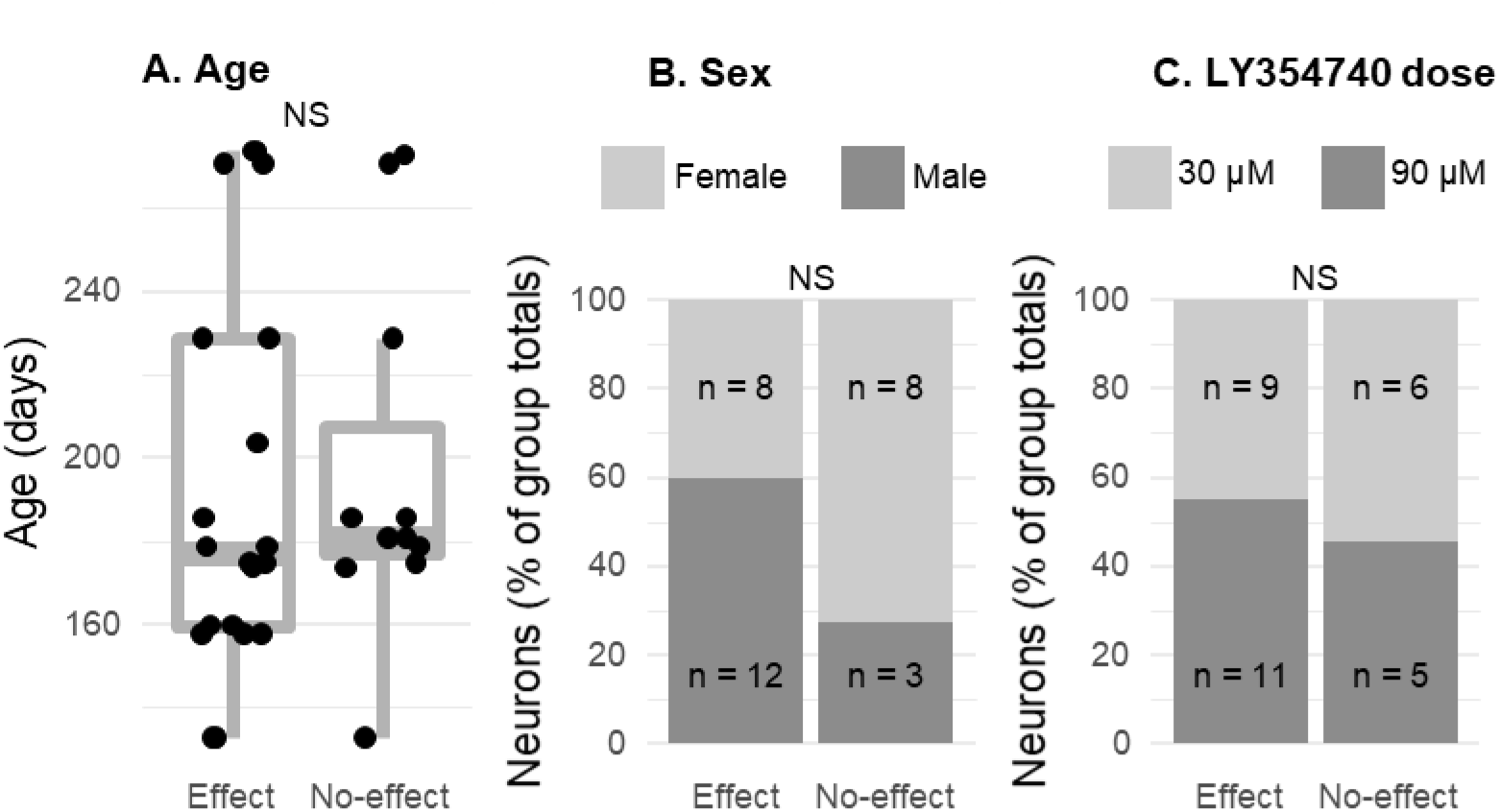
Association between the animals’ age, sex, drug dose and the effect of topical LY354740 application on rate-level functions. All neurons were split into two groups: those which had rate-level functions affected and unaffected by mGluR2/3 activation. No significant differences were found in age (**A**), sex (**B**), and drug dose (**C**) between these two neuron groups. NS p>0.05.

### The effect of mGluR2/3 activation on RLF changes is independent of agonist and drug application method used

The results described above were obtained using mGluR2/3 specific agonist LY354740 applied topically on the craniotomy above the IC. We performed experiments to further verify these results with two different positive control experiments. First, instead of topical drug application, we administered LY354740 using microiontophoresis (five neurons tested). This is a drug delivery method in which a small electric current is used to eject drugs from micropipettes attached to a recording electrode (Stone, 1985). This technique has been previously used to administer LY354740 (Copeland et al., 2012; 2013; 2017). Example RLFs recorded before and after iontophoretic LY354740 application are plotted in ***Figure 9A***. For a second positive control experiment we used topical drug application but utilized another highly specific group II mGluR agonist LY379268 (three neurons tested; Monn et al., 1999). ***Figure 9B*** shows example RLFs obtained before and after topical 30 μM LY379268 application. In both positive control experiments, mGluR2/3 activation resulted in similar effect on RLFs’ peak firing rates, slopes and thresholds as with our primary experimental protocol using topical LY354740 application. These data confirm that mGluR2/3 modulatory effect on sound processing in the mouse IC is robust and can be studied using different methods (further discussed in the Discussion). Negative control tests with topical and iontophoretic drug vehicle application did not result in RLFs’ modulation (six neurons tested).

**Figure 9.**
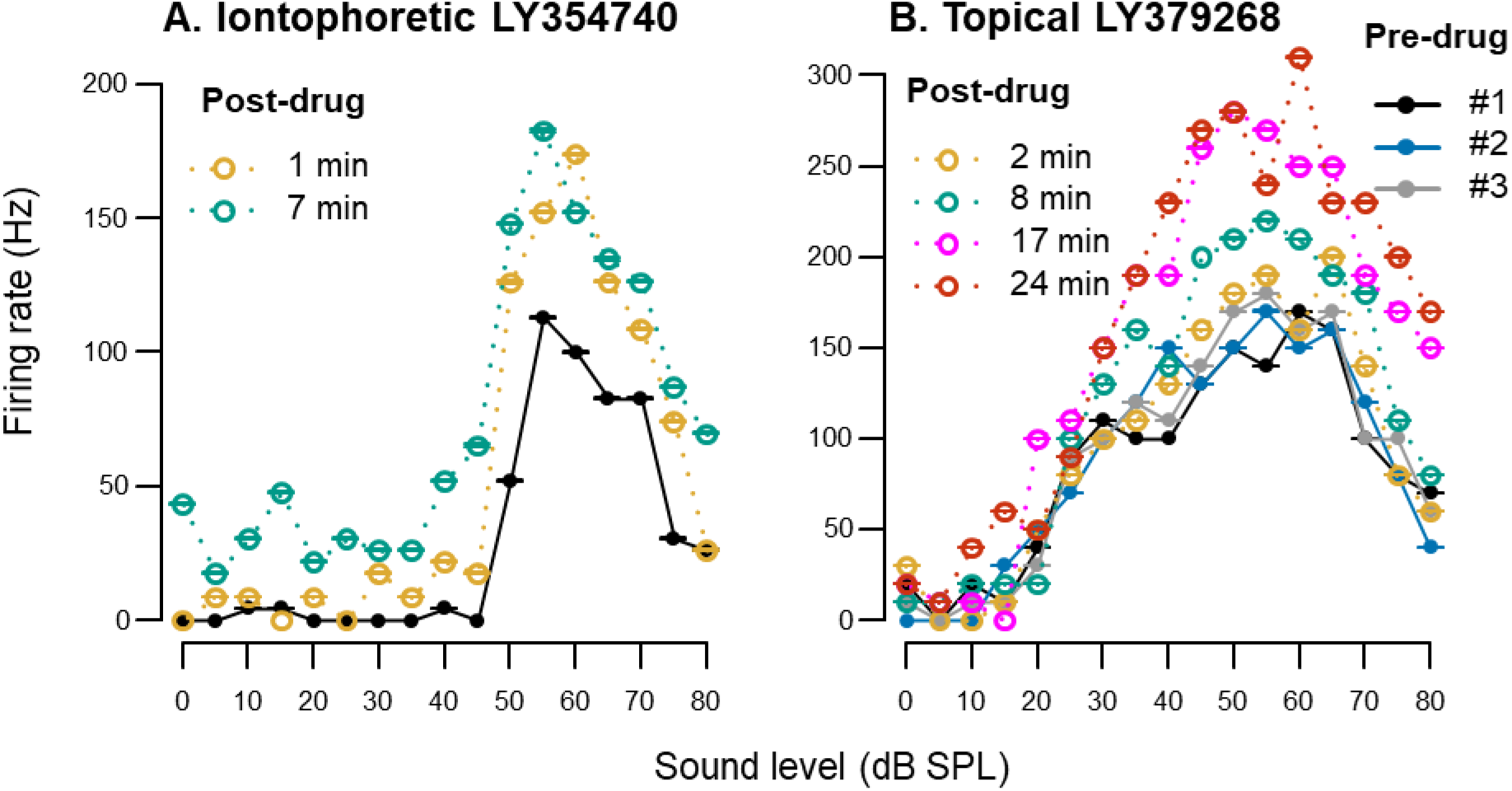
Examples of rate-level functions before and after mGluR2/3 activation: iontophoretic LY354740 ejection (**A**) and topical LY379268 application (**B**). Pre-drug rate-level functions are plotted in solid black, blue, and grey lines in both panels; the legend is only shown in panel **B**. Post-drug rate-level functions are color coded yellow, green, pink and orange to demonstrate drug effect over time; time after drug application is indicated for each individual rate-level function Error bars represent ± SEM of ten sound stimulus presentations at each sound level.

### MGluR2/3 activation increases spontaneous firing in the IC

In addition to sound-evoked activity, we also examined whether group II mGluR activation has effect on spontaneous firing. For this purpose, we studied the same 31 neurons described above for the RLF analyses. Thus, the same recordings were used for both sound-evoked and spontaneous firing analyses. Spontaneous firing rates were calculated from a 50-millisecond duration window prior to each tone presented during RLF data collection. For each RLF, a tone was presented 170 times: ten repetitions at seventeen sound intensity levels. Thus, spontaneous activity was assessed during the nine seconds period during each RLF test (170 trials * 50 ms = 9000 ms). As described above, several RLFs were obtained before drug application and several after, all of which were included in the analyses. Post-drug data collection typically lasted for about fifteen minutes (***Figure 7***).

Fifteen of 31 neurons were not spontaneously active before drug application, whereas after mGluR2/3 activation only six lacked spontaneous firing. In the entire sample, pre-drug spontaneous firing rates ranged from 0 to 5.5 spikes/second, while post-drug ranged 0 to 15.8 spikes/second. For statistical analyses, firing rates were rank-transformed because of non-normal distributions of raw data. Then, for each neuron, mean firing rate was calculated for all pre- and post-drug data and population-level analysis was performed using paired t-tests. In neurons assigned to the drug effect group (n = 20), there was a significant increase in spontaneous firing after drug application (t_19_ = 3.4, adj. p < .01; ***Figure 10A***). In contrast, no significant spontaneous firing change was found in the no drug effect group (n = 11; t_10_ = 1.9, adj. p = .09; ***Figure 10B***).

**Figure 10.**
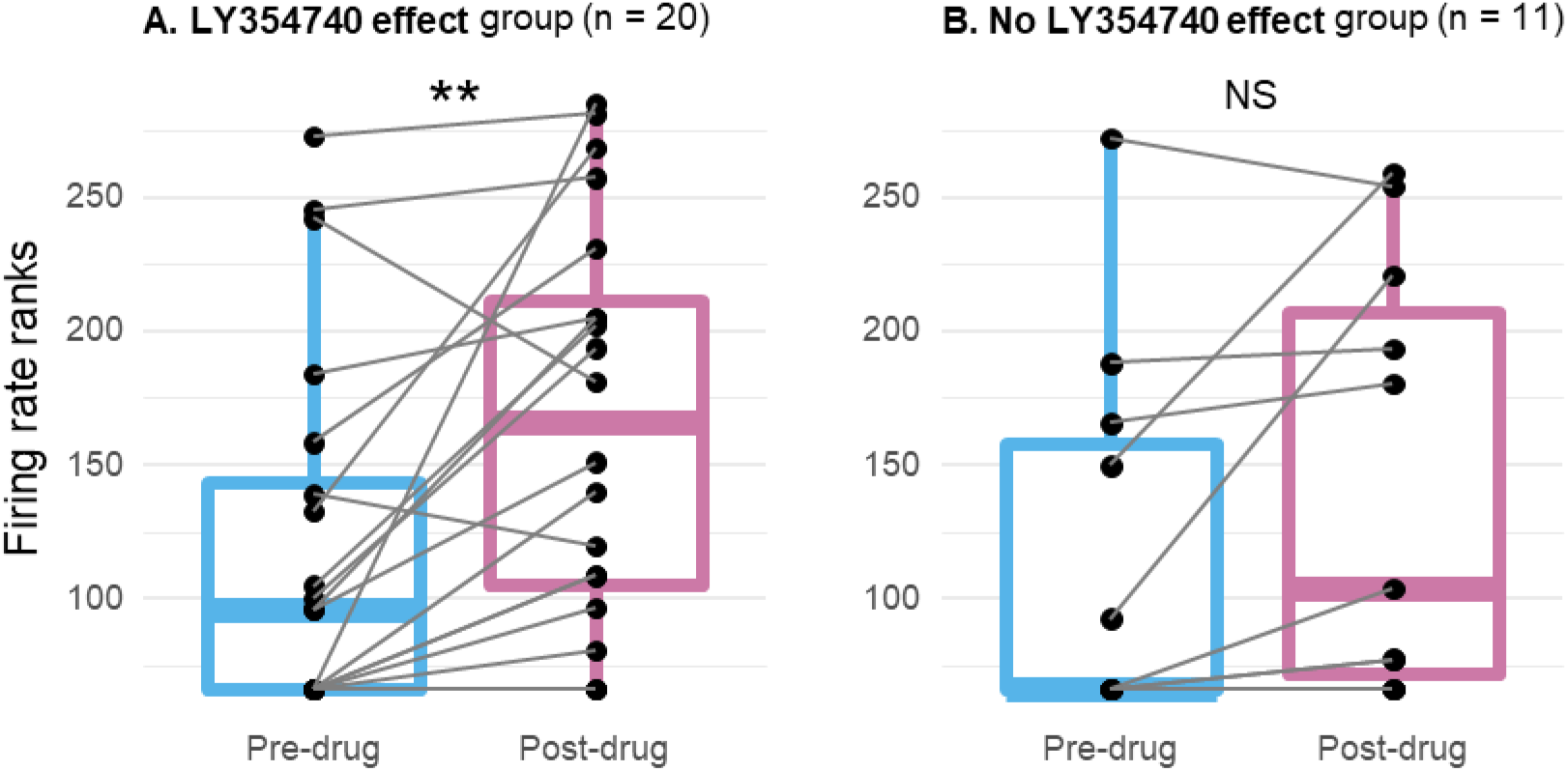
The effect of topical mGluR2/3 agonist LY354740 application on spontaneous firing rates. (**A**) Spontaneous activity significantly increased in neurons assigned to the drug effect group (n = 20). (**B**) Spontaneous activity did not significantly change in neurons assigned to the no drug effect group (n = 11). Each neuron’s pre-drug and post-drug means are connected by grey lines; population data is summarized with boxplots (the plot presents rank-transformed spontaneous firing rates). ** adj. p < 0.01, NS adj. p > 0.05.

### Neurons affected by mGluR2/3 activation were distributed throughout the IC area tested

At the end of the electrophysiological experiments, each recording site was labelled using iontophoretic dye ejection. From 31 neurons tested with topical LY354740 application, recording sites were successfully identified in twelve neurons. Fluoro-Gold labelling resulted in the largest proportion of identifiable sites (55% from 11 attempts), followed by Neurobiotin 350 (36% from 11 attempts) and Neurobiotin 488 (22% from 9 attempts). ***Figure 11*** shows examples of labelled recording sites. All dyes resulted in rather bright, well-confined labelling. This allowed for easy detection of recording sites in specific IC areas using a low (5X) magnification objective.

**Figure 11.**
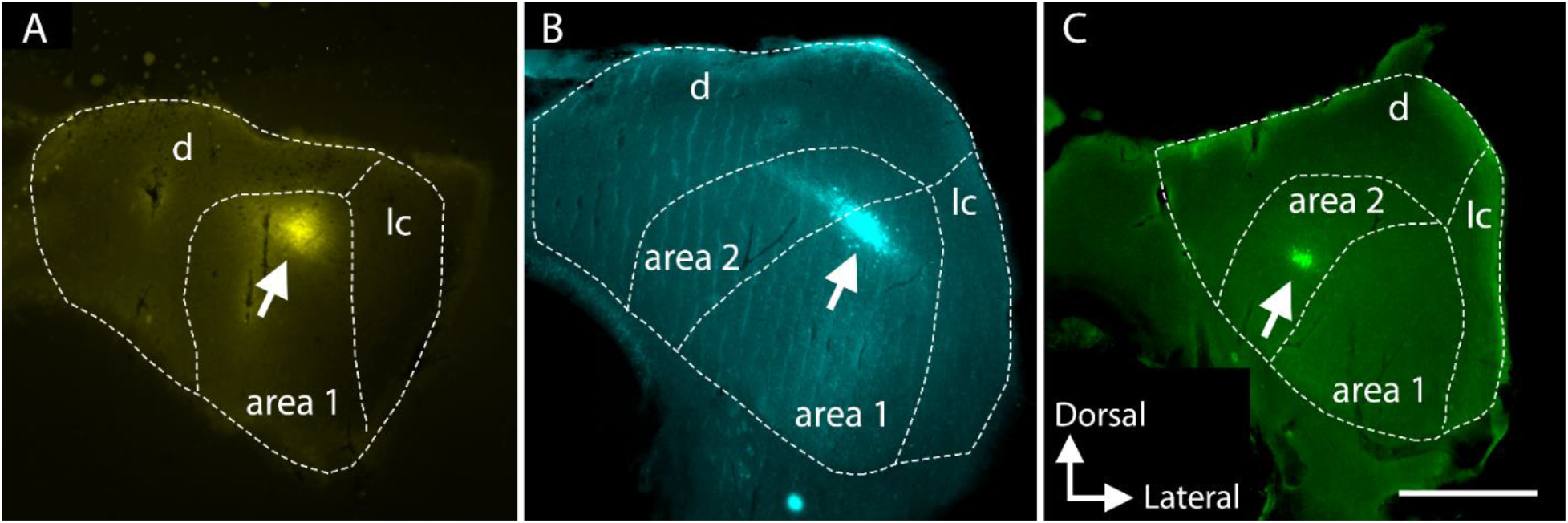
Example recording sites labelled with Fluoro-Gold (**A**), Neurobiotin 350 (**B**), and Neurobiotin 488 (**C**). Dashed lines indicate the boundaries of IC and its subregions: parts of the central IC nucleus - area 1 and area 2, as well as dorsal (d) and lateral (lc) cortex. Scale = 500 μm.

***Figure 12*** shows locations of successfully identified recording sites within the IC. Majority of recordings were from more dorsal IC parts due to concerns that the topically applied drug’s diffusion might be much lower in deeper regions. Indeed, drug effect was not observed in the two most deep successfully identified recording sites about 500 μm down from the IC surface. On the other hand, it is possible that drug effect occurred at such depths, but these recording sites were not recovered. In addition to histological recording site examination, all depths were noted during experiments based on how much the micromanipulator was vertically advanced into the IC. According to these data, the mean recording depth for neurons assigned to the drug effect group registered 251 μm (SD 107 μm), whereas the no effect group’s mean depth registered 333 μm (SD 134 μm). However, the mean difference was not statistically significant (t17 = 1.7, p = .10).

**Figure 12.**
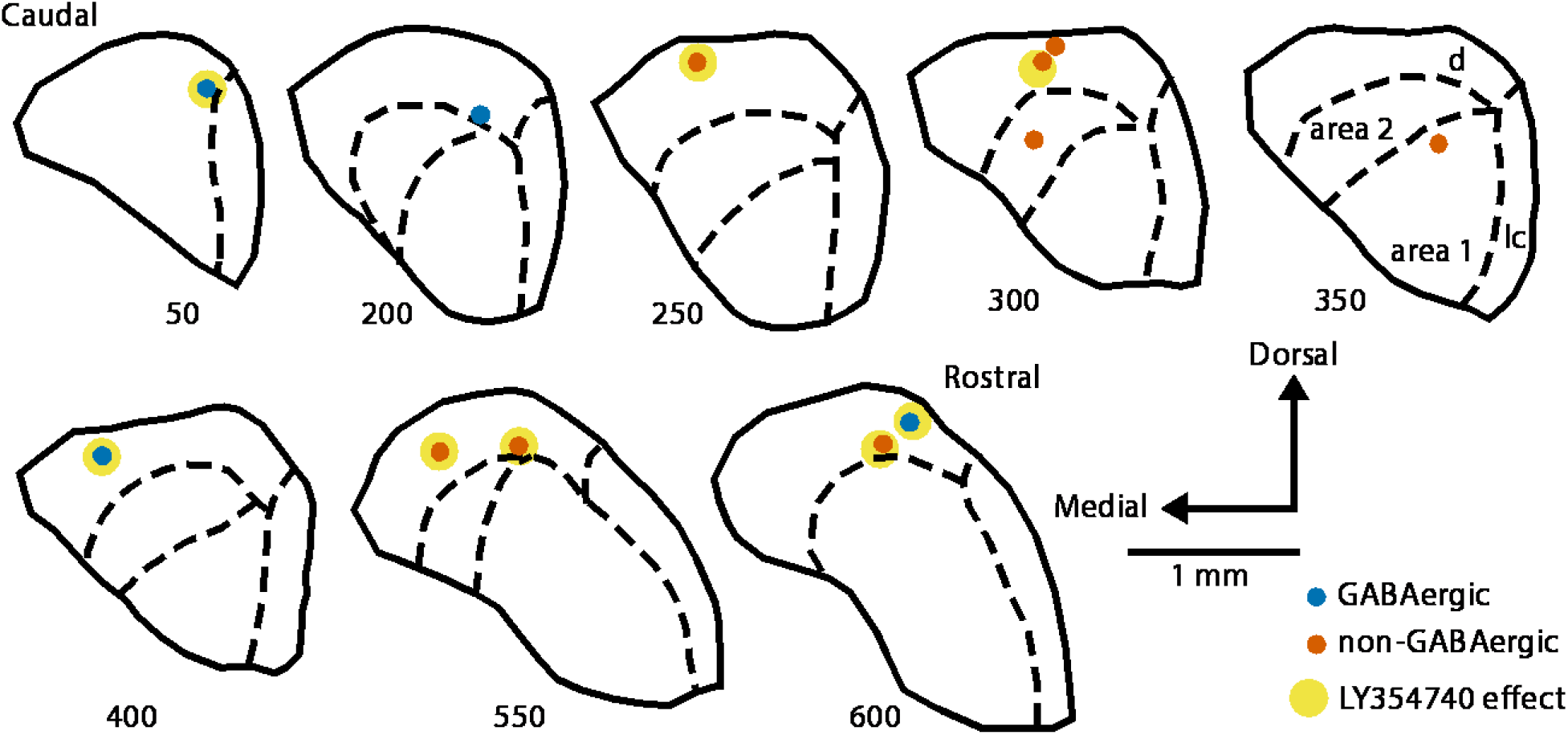
Successfully identified recording sites plotted on the schematic IC sections arranged from caudal to rostral. Numbers under each section indicate the approximate distance in μm from the caudal surface of the IC. Dashed lines indicate the boundaries of IC subregions: parts of the central IC nucleus - area 1 and area 2, as well as dorsal (d) and lateral (l) cortex (subdivisions are labelled only in the section at 350 μm from the most caudal IC part). GABAergic neurons are marked with blue dots, non-GABAergic with orange dots. The dots additionally marked with yellow indicate neurons assigned to the drug effect group.

Successfully identified recording sites were somewhat equally distributed along the rostral-caudal IC axis: six neurons in the more anterior half of the IC, the other six in the more posterior half (***Figure 12***). Five of six rostrally located neurons were from the drug effect group, whereas only three of six caudal neurons were affected by topical LY354740 application. Such result suggests a possibility that more rostral IC parts might be subjected to larger modulation by group II mGluRs compared to its caudal regions.

## Discussion

We utilized several experimental approaches—single cell electrophysiology, optogenetics, pharmacological and anatomical techniques—to investigate the role of group II mGluRs in the mouse inferior colliculus (IC). We first performed immunostaining and tested hearing thresholds to validate VGAT-ChR2 transgenic mice on a mixed C57BL/6J x CBA/CaJ genetic background. We confirmed that, as expected, hearing thresholds were normal and channelrhodopsin-2 was expressed in inhibitory cells. The use of transgenic animals allowed for optogenetic cell type identification and testing of a hypothesis that group II mGluRs modulate both excitatory and inhibitory cells. Recordings in the same neuron before and after pharmacological mGluR2/3 activation allowed us to show that after agonist application sound level processing is enhanced in a subset of both GABAergic and non-GABAergic IC neurons: rate-level functions became steeper, thresholds lower, and maximum firing rates higher. Spontaneous firing was also elevated after mGluR2/3 activation.

To the best of our knowledge, only one prior *in vivo* study was conducted to test the role of group II mGluRs in the IC (Voytenko and Galazyuk, 2011). Technical challenges related to recording the same neuron before and after receptor activation might discourage such projects. A more traditional approach is to deliver receptor agonists using iontophoresis, in which a small electrical current is used to eject a drug from a micropipette attached to the recording electrode. However, this method suffers from uncertainty regarding drug delivery *per se*, as well as unknown concentration of drug ejected into the tissue (Kirkpatrick and Wightman, 2016; Kirkpatrick et al., 2016). Drug concentration is important because even highly mGluR subtype-specific compounds can lose specificity at higher concentrations. Additionally, iontophoresis requires manufacturing multi-barrel electrodes and that is a complicated procedure. In our hands, commercially available multi-barrel electrodes (Kation Scientific) did not result in satisfactory signal-to-noise ratio. Furthermore, multi-barrel electrodes are significantly wider than single electrodes and inevitably cause more brain tissue damage, which might interfere with normal excitability. Considering these and other issues, we utilized topical drug application on the IC surface as our primary drug delivery method and performed iontophoresis to only verify the results. Although topical drug delivery approach is more commonly used in cortex (Happel et al., 2010; Self et al., 2014; Zhou et al., 2014; Rasmussen et al., 2019), there are several studies in which a drug was applied to the surface of the IC (e.g., Ji et al., 2001; Ji and Suga, 2009; Scott et al., 2018). In mice, the IC, like cortex, is conveniently located on the brain surface.

Topical drug delivery has several advantages. First, controlled drug concentration can be applied on the brain surface, minimizing the risk of losing the compound’s specificity for the receptors of interest. Second, a single recording electrode causes minimal brain damage and allows successful recording over the course of several days testing in the same animal. Additionally, signal-to-noise ratio was typically larger when a single electrode was used compared to multi-barrel, which was important for our goal to accurately compare the response of well-isolated neurons before and after drug application. A disadvantage of this approach is that the exact drug concentration at the recording site cannot be known. As mentioned above, the maximum is determined by the solution applied to the surface, but diffusion into the tissue will lead to a decrease in concentration to an unknown level. Nonetheless, we view topical drug administration as a valuable technique for testing the effects of pharmacological agents on IC physiology. Newly developing approaches that utilize light to activate and deactivate endogenous mGluRs may soon circumvent some of the problems of iontophoresis and topical application (e.g., Donthamsetti et al., 2019).

Our findings regarding the sound evoked activity—steeper RLF slopes, higher maximum firing rates, and lower thresholds—are partially congruent with the results from an *in vitro* study by Farazifard and Wu (2010). Both studies are in agreement that both GABAergic and non-GABAergic IC cell types are modulated by group II mGluRs. However, modulation direction differs. Farazifard and Wu (2010) showed reduced excitatory and inhibitory postsynaptic currents after mGluR2/3 activation, whereas our *in vivo* results demonstrated increased firing. The *in vitro* experiments were conducted in 8- to 17-days-old Long Evans rats, whereas our data was collected in 4- to 9-months-old mice. Age and species differences might explain the observed discrepancy. Indeed, it has been demonstrated that mGluRs are developmentally regulated in several brain regions (Catania et al., 1994; Defagot et al., 2002; Colantuoni et al., 2008; Hernandez et al., 2018); unfortunately, no data are available regarding the IC. Thus, future comparative biochemical and immunohistochemistry studies evaluating the effects of age and species on IC mGluR2/3 expression would be very valuable.

We found that spontaneous firing rates increased after group II mGluR activation. This result contrasts with results obtained using systemic intravenous mGluR2/3 agonist application: spontaneous firing was reported to be decreased (Galazyuk et al., 2019). As mGluRs have not been thoroughly studied throughout the auditory system and their expression patterns and physiological roles have not been described in detail, at this time it is difficult to propose a much more in depth explanation of the discrepancy other than stating the obvious—systemic drug application is vastly different from local application used in the present study. IC is often referred to as a hub of the auditory system because of converging ascending and descending auditory and some non-auditory inputs (Casseday et al., 2002; Gruters and Groh, 2012); systemically delivered drugs can bind to receptors in many brain areas and the effect on spontaneous firing could be relayed to the IC. When receptor activation is limited to the IC, the effect might be different due to the lack of impact on inputs from other regions. Therefore, to achieve a better understanding of how group II mGluRs might be utilized for possible treatment of tinnitus, which might be linked to elevated spontaneous activity (Sedley et al., 2019; Shore and Wu, 2019), it would be valuable to characterize the role these receptors play in spontaneous firing regulation not only in the IC but in other auditory structures as well.

In contrast to results reported here, an *in vivo* study by Voytenko and Galazyuk (2011) found no effect of iontophoretically delivered group II agonist on either sound evoked or spontaneous firing in the IC. The animals in that study were 2.5 months younger than the youngest mice used in the present study. None of the recording sites were labelled in the 2011 publication. It is possible these data were obtained from deeper IC areas which might have lower/no mGluR2/3 expression. It is likely that only a specific subset of cells expresses these receptors; a small sample size in a report by Voytenko and Galazyuk (2011) might have contributed to the lack of observed effect. Therefore, it would be worthwhile to investigate possible differential receptor expression between IC subdivisions. Information about receptor density would also provide some clarification regarding the lack of drug effect in a subset of neurons described in the present study.

Findings in line with those reported here—increased sound-evoked and spontaneous firing after group II mGluR activation—were described in other brain areas. Increased spontaneous excitability was demonstrated *in vitro* in hippocampal CA3 pyramidal cells (Ster et al., 2011) and *in vivo* in the cochlear nucleus (Sanes et al., 1998). An *in vivo* study by Jin et al. (2017) showed that iontophoretically applied mGluR2/3 agonist increased Delay cell firing in the primate layer III dorsolateral prefrontal cortex. Both Jin et al. (2017) and Ster et al. (2011) examined subcellular mGluR2/3 localization and found evidence for both pre- and post-synaptic sites, confirming earlier findings (Petralia et al., 1996; Tamaru, 2001). Traditionally, pre-synaptic receptors were the focus of most research. Related is the fact that group II mGluRs are coupled to Gi/Go proteins which are associated with inhibition of the adenylyl cyclase and cyclic adenosine monophosphate (cAMP) formation (Conn and Pin, 1997), leading to reduced glutamate levels in the synapse and subsequently decreased neuronal firing. In light of these findings and the findings of the present study, it is possible enhanced firing may be exclusive to or more pronounced in brain regions containing primarily post-synaptic mGluRs2/3. For example, Tyszkiewicz et al. (2004) demonstrated that group II mGluR activation increased post-synaptic glutamate ionotropic NMDA receptor currents in dissociated rat prefrontal cortex pyramidal neurons. As we learn more about receptor subcellular localization, and expand efforts to disambiguate subtype specific roles (mGluR2 versus mGluR3; Copeland et al., 2012; Jin et al., 2018; Wood et al., 2018), it should be possible to better explain how group II mGluRs can exert both inhibitory and excitatory actions. It remains to be tested whether in the mouse IC post-synaptic receptors may be a major mGluR2/3 component.

What is the biological relevance of enhanced sound-evoked and spontaneous firing in the IC following activation of group II mGluRs? In the hippocampus, group II mGluR activation increased spontaneous neuronal excitability which resulted in synchronous network activity in the theta range (Ster et al., 2011). Although the exact function of hippocampal theta oscillations is not known, this might be vital for optimal memory encoding (Leung and Law, 2020). In the auditory system, however, synchrony and increased spontaneous activity have been investigated as plausible mechanisms underlying tinnitus (Shore et al., 2016; Shore and Wu, 2019) and audiogenic seizures (Garcia-Cairasco, 2002). Thus, it seems that elevation of spontaneous firing by activation of mGluR2/3 receptors in the IC could have negative consequences. Still, could elevated spontaneous firing have a positive effect on sound processing? Under physiological conditions, when sound level increases, glutamate release becomes more substantial and clearance of glutamate by uptake mechanisms is delayed (Scanziani et al., 1997). This results in increased glutamate concentration in and around the synapse and more mGluRs are activated, possibly including receptors at the peri-synaptic and extra-synaptic membranes (Jin et al., 2017; 2018). Thus, mGluRs2/3 may enhance sound-level processing primarily at higher sound intensities. In our study, however, the exogenous agonist application must have been uniform across sound levels and increased firing irrespective of sound intensity.

Many interesting questions remain to be answered regarding a role of group II mGluRs in the IC. We hope our results will encourage further investigations by providing critical *in vivo* data that group II mGluRs enhance both sound-evoked and spontaneous firing in the mouse IC in both GABAergic and non-GABAergic cell types.

## Acknowledgements

Sincere special thanks to Dr. Emily Hazlett for helping to set up the electrophysiological experiments using the BrainWave software, Dr. Douglas Oliver for advice on optogenetics and other generous suggestions, Dr. Donald Caspary for assistance with iontophoresis, Dr. Calvin Wu for recommendation regarding the RLF fits, and Dr. Yong Lu for comments on an earlier draft of the manuscript. This work was supported by National Institutes of Health Grants R01 DC016918 (to AVG) and R01 DC004391 (to BRS).

